# Predicting an epistasis-rich genotype-phenotype map with a coarse-grained bottom-up model of budding yeast polarity

**DOI:** 10.1101/2020.11.30.403758

**Authors:** Werner Karl-Gustav Daalman, Liedewij Laan

**Affiliations:** Department of Bionanoscience, TU Delft, Delft, the Netherlands

## Abstract

Accurate phenotype prediction based on genotypical information has numerous societal applications, such as design of useful crops of cellular factories. However, the prevalence of epistasis, a phenomenon that prevents many biological systems to perform in accordance with the sum of its parts, necessitates modelling the complex path between genotype and phenotype. Defining intermediate levels in this path reduces the complexity of prediction, and may also elucidate the phenotype coupling to other levels by evolution. Inconveniently, the latter requires definitions that maintain biophysical justification from the bottom-up, which conflicts with tractability. By means of a cell growth model, we exemplify a resolution for this conflict by polarization of Cdc42p in budding yeast, a process requiring clustering of active Cdc42p to one zone on the membrane and known to generate ample epistasis. Concretely, our model parsimoniously encompasses constant membrane area growth, stochastic Cdc42p turnover and a simple, justifiable polarity rule we define as the ‘mesotype’. Through intuitively interpretable simulations, we describe previously documented, yet puzzling epistasis inside the polarity module. Moreover, we generate evolutionary relevant predictions e.g., on environmental perturbations, which are general enough to apply to other systems. We quantify how poor growth medium can equalize fitness differentials and enables, otherwise very distinct, evolutionary paths. For example, the fitness of the crippled Δ*bem1* relative to WT can easily be raised from 0.2 to above 0.95. Finally, we can extend our predictions on epistasis to other modules. We determine that modelled epistasis predictions only add predictive value when functional information of the involved modules is included. This inspires a road-map towards modelling the bidirectional genotype-phenotype map for other model systems with abundant interactions, where the intermediate levels reveal targets that evolution can optimize and facilitate a biophysical justifiable incorporation of epistasis.

**Author summary:** Efforts to understand how traits follow from genes facilitate a broad range of applications. For example, crops can be engineered faster to better resist drought, salt and heat stress, and medicines can be better tailored to individuals. Unfortunately, the path from genes to traits can generally involve a complex interplay of hundreds of genes and gene products whose individual contributions can be heavily context-dependent. In this work, we provide the proof-of-concept in a relatively simple system for a road-map towards elucidating this path. We have constructed a cell growth model for budding yeast, only involving simple rules on membrane growth, protein production and centrally, polarity, the process where yeast chooses the future division site. Despite the simplicity, the polarity rule is fully justifiable from underlying biophysics. Model simulations show good accordance with formerly puzzling traits, and also predict the ease with which the environment can change evolutionary paths. While lab conditions may prohibit the emergence of certain polarity mutations, this becomes much more feasible ‘in the wild’. The tractable model nature allows us to extrapolate the context dependence of mutational effects beyond polarity, showing that this method for understanding trait generation also helps to elucidate protein evolution.

## Introduction

Many fields, such as personalized medicine [1], agriculture [2], chemical production [3] and forensics [4], will greatly benefit from advances in understanding of the so-called genotype-phenotype (GP) map, the way that traits are connected to genes. However, this connection can be quite complex even for known heritable traits (“missing heritability”) [5], limiting the power of genome-wide association studies [6]. On the one hand, one gene can be responsible for multiple traits, pleiotropy, although this may not always be very common [7]. On the other hand, multiple genes can contribute to one trait. Frequently, their individual effects are non-additive in humans [8,9], but also in model systems as *Escherichia coli* [10] or *Saccharomyces cerevisiae* (budding yeast) [11], a phenomenon known as epistasis. While some molecular mechanisms for epistasis are known [12] which can be inconsequential for fitness evolution [13], the presence of epistasis complicates the predictions of phenotypes from genotypes and consequently gene evolution [14,15]. Therefore, predictions on epistasis constitute an important challenge for GP-map models.

As a modelling tool to more easily decompose the GP-map, intermediate levels can be defined as stepping stones [16], which can be brought under the general denominator of causally cohesive genotype-phenotype models [17]. An intermediate level may provide an entry point for additional observables that fine-tune predictions, but an abstract, unobservable entity as a definition is also possible. Most importantly, a level serves to break up and re-bundle the intertwined paths from individual genes to traits such that a more modular and hence more tractable picture arises. In that view, a suitable level definition acts as a tree which branches out to otherwise difficult to connect genotypes and phenotypes.

Multiple level examples exist, such as the biofunctional gene ontology level (ontotypes) [18], the network based trophic level [19], the diffuse endophenotypes [20] and the mathematical system design space [21]. Ideally, a one-level-fits-all approach exists, where the level definition facilitates understanding of the emergence of phenotypes from genotypes, while at the same time elucidating the handles for evolution, the reverse path in the GP-map. This requires steering away from phenomenological or statistical formulations to move towards biophysically sound versions, while at the same time maintaining tractability which often complicates bottom-up approaches. Consequently, the generation of a suitable level definition for the tractable bidirectional path in the GP-map, if possible, involves coarse-graining of the underlying biophysics, specifically of molecular interactions and protein transport.

A promising attempt is the iMeGroCy model [22], where growth and cell cycle processes are simplified, and more details are kept for the module of interest, in this case the metabolism, which follows Michaelis-Menten kinetics. While effective in modularizing e.g., the pedigree phenotype emergence as a function of medium in *S. cerevisiae*, it is not straightforward to extrapolate this approach to other modules. The kinetics crucially assume well-mixed components, neglecting spatial heterogeneity arising from crowding in the cell [23]. Furthermore, coupling of diffusion and reactions involves conditions for pattern formation [24] that should be taken into account. Inclusion of spatio-temporal information is also known to be essential to understand the evolution of a network [25]. We therefore construct a cell growth model that encompasses the lessons derived from rigorous reaction-diffusion analysis, and maintains the simplicity across other growth features. The level definition associated with coarse-graining the molecular details will be called the ‘mesotype’.

For the purpose of testing this bottom-up modelling approach, we require a model system known to exhibit ample epistasis (e.g., in doubling times) [26], namely polarity establishment in *S. cerevisiae*. Here, the unicellular organism budding yeast breaks its internal spherical symmetry to direct bud growth in one direction. This process is essential for the proliferation of the cell and relies on correct functioning of Cdc42p [27]. The mechanics behind this process is known to large detail (except the role of unexpected wildcard Nrp1p [26]), and involves clustering of the active form of small GTPase Cdc42p, which is bound to a GTP molecule, to one patch on the plasma membrane (Fig. 1A). In a wild-type (WT) background, rapid Cdc42p clustering is governed by a positive feedback involving Bem1p and Cdc24p [28], the relevant guanine nucleotide exchange factor (GEF) for Cdc42p [29], which appropriately transport and activate Cdc42p. By contrast, its deactivation outside the membrane patch is ensured by GTPase activating proteins (GAPs), a protein class to which Bem2p, Bem3p [29], Rga1p [30] and Rga2p [31] pertain.

**Fig 1.**
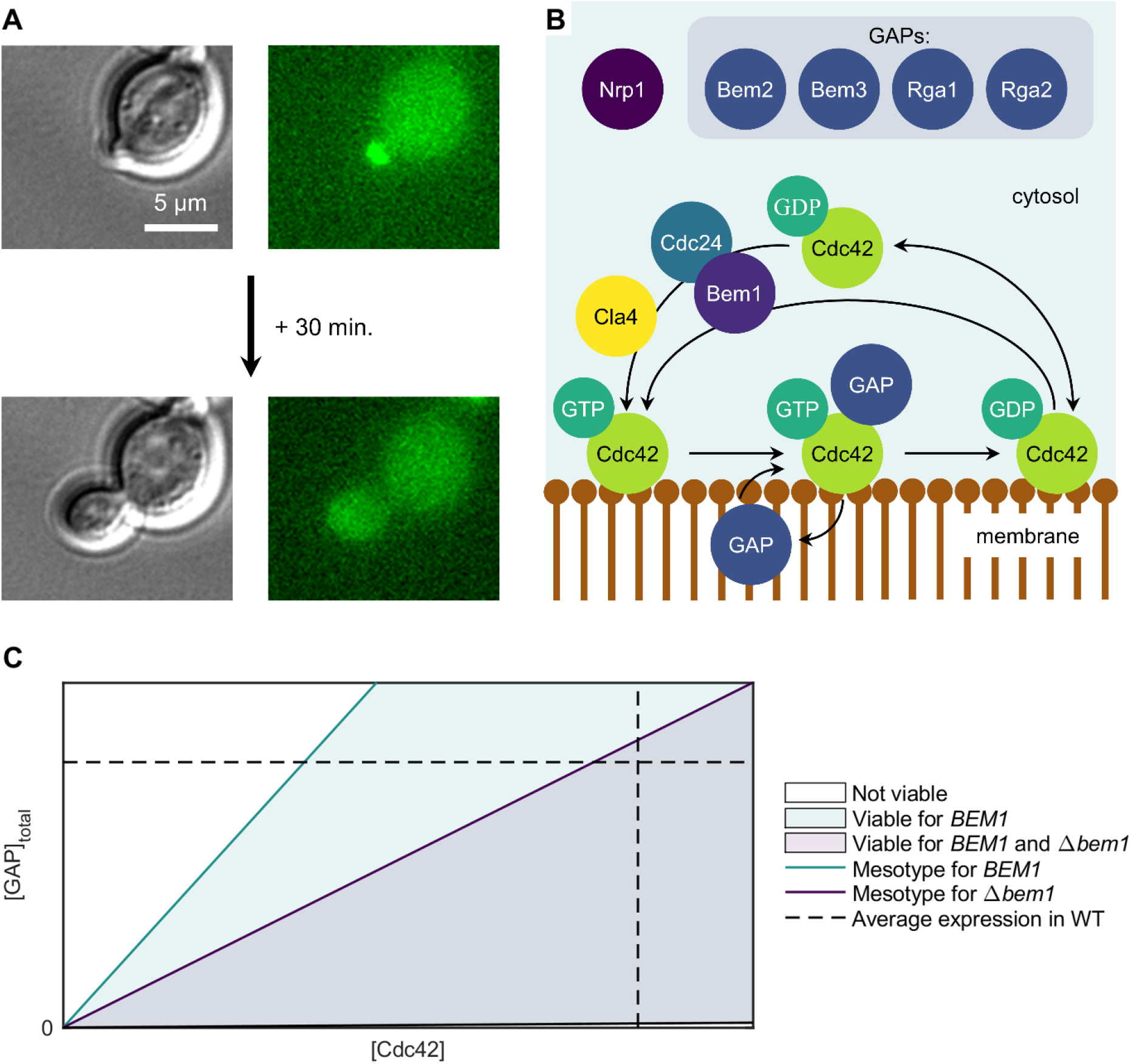
Yeast polarity as suitable genotype-phenotype map model for epistasis description. (A) Brightfield (left) and widefield fluorescence example images of a polarizing budding yeast cell (scale bar 5 μm). Key for polarity is clustering of active Cdc42p (of which a binding partner is fluorescently labelled in the images) to one zone on the membrane. This location marks the site of polarized growth. (B) Schematic overview of polarity protein core (proteins not to scale). 6

A positive feedback for (active) Cdc42p-GTP is mediated by either the Bem1p-Cdc24p complex, and likely to lesser extent by Cla4p. For Nrp1p it is unclear how it mechanistically links to other components. (C) Schematic diagram depicting phenotype (viability) as function of genotype through Cdc42p (active and inactive) and GAP concentration in the cell with or without Bem1p. An intermediate, the ‘mesotype’, is defined here as the limiting Cdc42p concentration. Epistasis is readily observed as the same increase in e.g., GAP concentration can yield inviability in the Δ*bem1* background but not in the *BEM1* background.

In absence of Bem1p, GAPs can more easily deactivate even the Cdc42p localized in the main patch that marks the future division site, which would generate a lethal situation for the cell. This can be circumvented if the abundance of Cdc42p is large enough to continuously sequester the GAPs found around the main patch, forming a rescue mechanism to establish polarity when combined with a generic positive feedback [32], such as through Cla4p [33] (Fig. 1B). Theoretical analysis of the underlying reaction-diffusion equations reveals a strong dependence of the ability to polarize success on the GAP/Cdc42p copy number ratio, where a broader range is viable in the presence of Bem1p [32] (Fig. 1C). This motivates a coarse-graining of the protein dynamics to a threshold for the protein concentration, which forms the mesotype level definition in this context.

## Results

### Coarse-grained bottom-up model design

#### Coarse description of cell expansion

We modeled the yeast cell cycle as a process involving three modules, namely coarse-grained cell growth, protein turnover and cell polarity (Fig. 2A). A parsimonious approach to cell membrane growth was chosen consisting of two stages of constant membrane area growth, as several alternative formulations proved immaterial for phenotype description (S1 Fig.). The membrane expansion rates (*C*_*1*_ for G1, *C*_*2*_ elsewhere) were brought in decent agreement with literature [34,35] (see S2 Table for further justification of parameter choices.).

**Fig 2.**
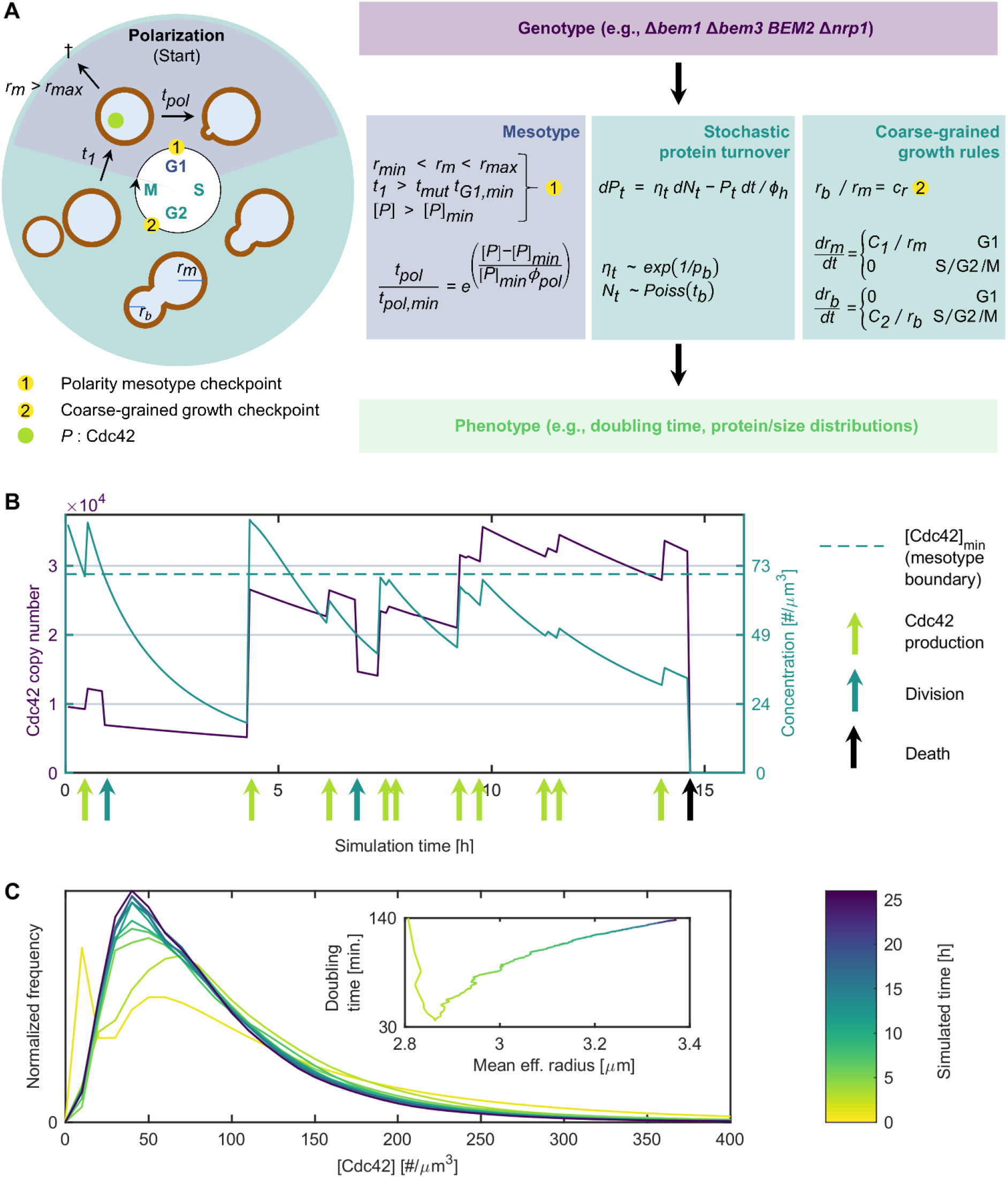
Coarse-grained, bottom-up growth model integrating the polarity mesotype to facilitate epistasis and phenotype prediction. (A) Schematic depiction of the translation of the budding yeast cell cycle to model processes and parameters. Central is the moment of polarization which occurs when the cell has sufficient size, has been sufficiently long in G1 and has a Cdc42 concentration (abbreviated as [*P*]) exceeding a threshold, the latter defining the mesotype checkpoint for this system. Together with a coarse description of cell growth (constant membrane expansion of either mother or bud and a bud size checkpoint), stochastic protein production and deterministic degradation, this allows construction of the genotype-phenotype map. (B) Example trace of the Cdc42 copy number (blue) and concentration (green) of a single cell, which are subject to protein production and degradation (and dilution for concentrations). The cell must exceed the mesotype threshold ([*P*]_min_) before division can take place. When this is delayed for too long, the cells expands beyond *r*_*max*_ and the cell dies (after almost 15h). (C) Convergence of Cdc42 copy number distribution during simulations. Simulated time since ancestor is approximate as birth times of the cells in the starting population are distributed across an 83 min. bandwidth. The inset shows how the estimates of the population doubling time and the average effective cell size equilibrate as a function of time.

The first stage involves isotropic growth of a spherical cell of radius *r*_*m*_*(t)*, which corresponds to the G1 phase including the Start transition, at the end of which a checkpoint must be passed, which is further explained in the following paragraph. Thereafter, the cell switches to the second stage of growth, where the membrane grows in a polarized manner defining a bud with radius *r*_*b*_*(t)*, while the rest of the mother cell retains its size. The bud membrane growth area is constant for the modelled equivalent of the S, G2 and M phase but larger than for the mother in G1. Bud growth lasts until the second checkpoint, at which the bud proceeds as an independent cell when it has reached a sufficient size (*r*_*b*_=*r*_*m*_ *c*_*r*_).

#### Biophysically justifiable mesotype inclusion

The cells grow isotropically until three conditions are met, defining the first checkpoint. Firstly, the radius *r*_*m*_ of the cell must exceed the minimum size threshold *r*_*min*_. Secondly, the time in this stage (*t*_*1*_) exceeds a minimum time (*t*_*G1,min*_), which may be modified by a factor *t*_*mut*_ for certain mutations with respect to WT. These two criteria result from key events in the timing pathway, particularly cell size dependent control of Cln3p arrival to the nucleus by Ydj1p [36]. Finally, the concentration of Cdc42p, [*P*], must exceed a minimum concentration threshold [*P*]_min_. The existence of the Cdc42p concentration threshold, which we define as the ‘mesotype’ for a particular mutant, follows from rigorous theoretical and experimental analysis of reaction-diffusion equations of the polarity network [32].

Once all three conditions are met, isotropic growth continues for a period of *t*_*pol*_, which lasts at least *t*_*pol,min*_ and depends exponentially on the relative excess Cdc42 concentration above the threshold (scaled by *ϕ*_*pol*_). This is a simplified representation of the results in [32] and reflects the period where Cdc42p clusters to one zone in the membrane. As it can occur that the Cdc42 threshold is never exceeded while growth continues and concentration are diluted, the cell is considered dead when its radius exceeds the maximum size *r*_*max*_.

#### Noisy protein production

Whether the Cdc42 concentration condition is met depends also strongly on protein production, which is modelled as a stochastic process. Since mRNA lifetime of Cdc42 is much smaller than its protein half-life *t*_*h*_ [37,38], Cdc42p production essentially follows from instantaneous bursts, which are modelled as an compound Poisson process burst process *N*_*t*_ with exponentially distributed size *η*_*t*_ (on average *p*_*b*_) at exponentially distributed intervals (on average *t*_*b*_) [39]. Because it may be important for the precise crossing time of the polarity threshold, we avoid absorbing protein degradation in an effective burst size, by explicitly adding degradation to the Cdc42 copy number process *P*_*t*_. Stochasticity in total GAP copy number is not included as the cell-to-cell variability is much less than for Cdc42p (GAP coefficient of variation < 0.15, only just above the smallest measured value of 0.10 [40], compared to 0.83 for Cdc42p measured in this study).

### Coarse-grained bottom-up model verification and validation

Firstly, the model design was verified by simulations of the computational model implementation (see Materials and Methods) which allowed tracking the states of individual cells or the population. Fig. 2B shows a Cdc42p copy number and concentration time trace of a single cell. The Cdc42p production is burst-like and occurs as indicated on the time axis. The copy number trace shows the proteins degrade between these bursts, and there is also dilution due to cell volume growth for the concentration curve. In this example, divisions occur twice shortly after checkpoint 1 has been passed, which also implies exceeding the mesotype concentration threshold. Ultimately, this cell fails to exceed the threshold a third time and dies after exceeding the maximum size *r*_*max*_, as designed. Fig. 2C shows the rate of convergence of relevant population phenotypes (without plotting the dilution step). After the population has grown approximately 25 hours counted from the ancestor seed, the Cdc42p distribution has largely converged. The size and doubling time change 0.6% and 0.9% respectively across the last 200 minutes, well within the typical experimental error (see S2 Fig. for an example of results including dilution).

Secondly, we turned to the model validation, where five parameters (four mesotype thresholds, and one *nrp1*-dependent G1 time factor *t*_*mut*_) are fitted. Non-trivial previously measured observables are considered mostly from [26]; strong epistasis in growth rates between GAP mutants only in the Δ*bem1* background, strong epistasis between *BEM1* and *NRP1*, and non-monotonous optimization of G1 times for (reconstructed) experimentally evolved mutants starting from Δ*bem1*. For the latter phenotype, the acceleration of G1 speed of the Δ*bem1* cells, despite its poor fitness, compared to WT cells is particularly noteworthy. This combines to a total of 20 phenotypes, well identifying the five free parameters.

#### Description of the coarse GP-map exhibiting epistasis

The simulated growth rates of polarity mutants of [26] were calculated as a function of mesotype ([Cdc42]_min_), which scales linearly with total GAP concentration [GAP]_*tot*_ due to the cone-like structure of Fig. 1C. These can be converted to relative fitness values through division by the WT growth rate. Fitness values in presence of *NRP1* were brought in accurate accordance with experiments of [26] (Fig. 3A) and consequently, the observed GAP epistasis is well (and robustly, see S1 Fig.) described. The *nrp1* background was not always well fitted (5/8 correct within experimental error), although these mutants suffered from relatively large experimental uncertainties. The four fitted mesotype threshold concentrations are consistent with the *bem3* deletion effect that is twice as large as for *bem2*. Given the GAP abundancies [41], this sets the *in vivo* Bem3p effective GAP activity to be almost four times as large as for Bem2p, a difference much less pronounced than measured *in vitro* [29].

**Fig 3.**
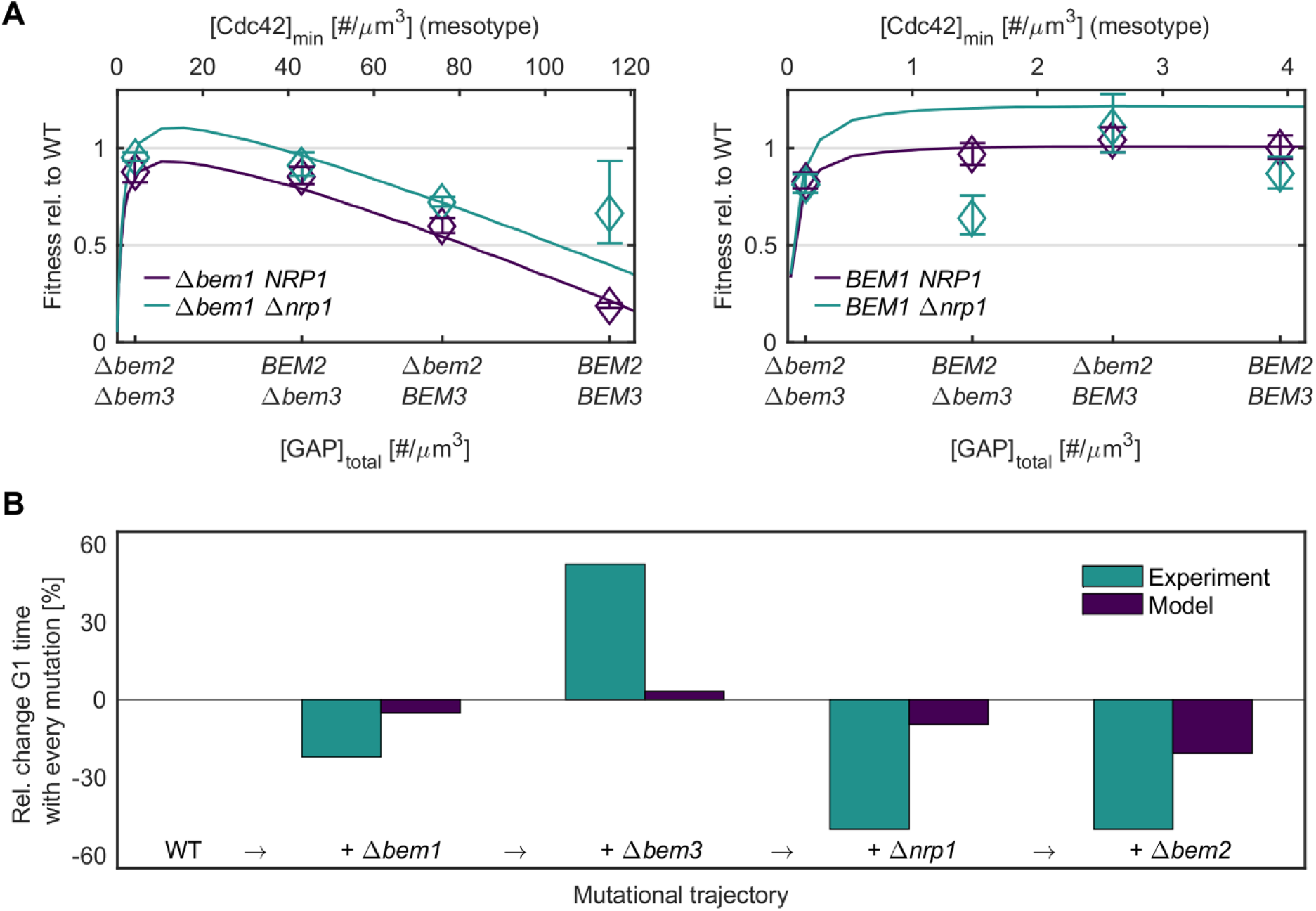
Comparison of the coarse-grained growth model to experimental data on coarse and subtler GP-maps. (A) Experimental fitness values relative to WT (phenotypes) for 16 different polarity genotypes [26] denoted by diamonds, which are fitted by the model as depicted by the dark purple and emerald lines for *nrp1* and *NRP1* background respectively. GAP genotypes can be linearly linked to the minimum Cdc42 concentration to polarize, the mesotype, as displayed on the top horizontal axis. Error of the *BEM2 NRP1* was not available and conservatively guessed. (B) The detailed phenotype of minimum G1 time as displayed by WT and four polarity mutants, comprising a single evolutionary trajectory. Experimental values are from [26] in emerald (defined there as tie to first polarity spot), model values are in dark purple (defined here as the time in G1 until both the size and time criteria are met). Both cases are normalized to their respective WT values, such that each column denoted the relative change in G1 time compared to the previous step in the trajectory.

#### Incorporation of a subtler GP-map

While doubling times represent a rather coarse phenotype, an example of the more detailed traits that can be modelled is time spent in G1. To this end, simulations were performed with half the normal membrane area rates *C*_*1*_ and *C*_*2*_, to mimic the poorer content of synthetic medium in which experiments from literature [26] were performed. The observed trends in G1 times along the evolutionary trajectory from WT to the fully evolved mutant in that paper were qualitatively matched, including the unusual inversion for the Δ*bem1* (Fig. 3B). The logic behind this inversion is that for WT cells in slower growth medium, the size requirement is the most important criterion for the first checkpoint of Fig. 2A, which can last longer than the minimum G1 time. By contrast, the on average less fit and larger Δ*bem1* cells are relatively more stalled by the minimum time criterion, and the long overall cycle times arise due to lengthy other phases.

A more realistic (and less coarse-grained) modification of the modelled cell cycle progression can improve the quantitative match. Suppose for example the *Δbem1* cells if the assumed minimum G1 time set is not a constant but a distribution (times for symmetry breaking in daughter cells can be quite stochastic [42]). Some Δ*bem1* cells have an early opportunity to fulfil the mesotype threshold concentration requirement, with which they usually struggle, while others are delayed more. This increases the cell-to-cell variation in fates in G1, since cells with fast G1 times are most likely to generate a first spot, while slow cells never generate this spot do not show up in the statistics. This is how less coarse-graining can lead to a larger decrease in G1 times than is the case with constant *t*_*G1,min*_.

### Genetic interaction predictions

#### Poorer medium quality reduces fitness differentials

As aforementioned, the effect of the environmental effects such as changes in growth media quality can be integrated in the model through a change in membrane area growth rates *C*_*1*_ and *C*_*2*_. To assess the evolutionary consequences of poorer medium content, we considered a roughly three-fold area growth rate range that caused WT fitness to span between 0.5 and 1 (normalized to maximum growth). Fig. 4 shows the fitness ratio for various media within this range between the Δ*bem1* and *BEM1* background, as a function of GAP concentration, visualizing the trend of smaller fitness differentials for decreasing GAP concentrations and decreasing medium quality.

**Fig 4.**
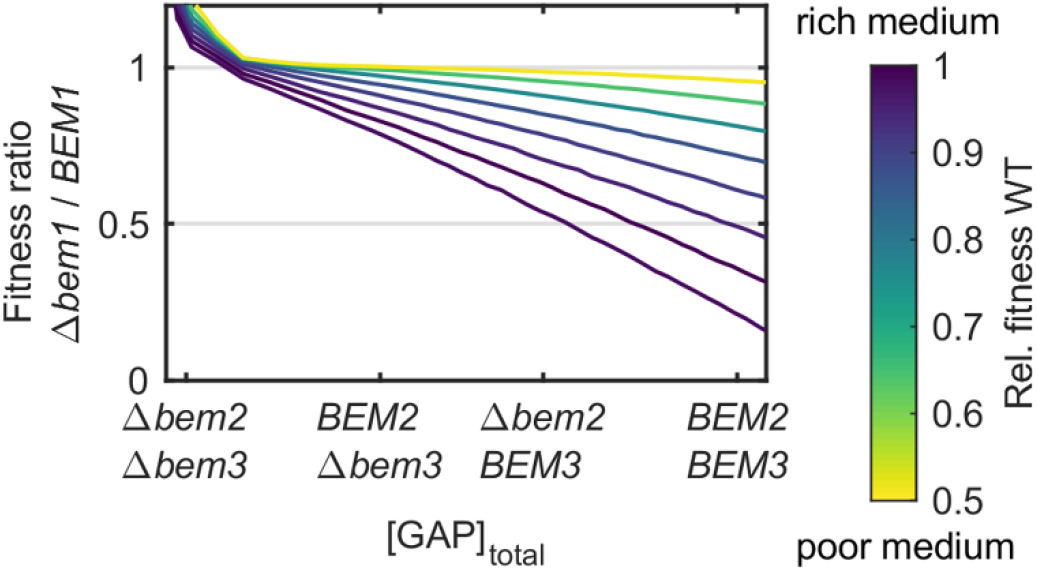
Growth model predictions of the environmental effect on polarity epistasis. Simulated fitness differences between *BEM1* and *bem1* backgrounds as a function of medium quality, which is integrated in the model through varying cell membrane area growth rates. The colors depict these rates through their associated WT doubling times. Generally, poorer medium reduces differences in fitness and genetic interactions between GAPs when comparing the *BEM1* and *bem1* backgrounds.

The intuition for this result is as follows. As seen in Fig. 3A, the Δ*bem1* background suffers from the high Cdc42p concentration threshold, relevant at checkpoint 1 (fig. 2A), and recover fitness when this threshold is lowered by successive GAP deletions. Fig. 2B had in turn shown the strong negative influence of dilution on the ability to exceed this threshold. Therefore, *Δbem1* cells benefit greatly from reducing the speed of membrane growth, while WT cells, for which the threshold is not a problem at all, only suffer from slowing down the membrane growth. An unmodelled inhibitor of this effect would be a reduced Cdc42p production in medium with lower quality. However, Cdc42p expression is at least known to remain stable upon switching from dextrose to ethanol, an inferior carbon source [43].

#### Information on biological function of mutated genes are a prerequisite for predicting epistasis

To assess whether we can extend the model predictions beyond polarity, we focus on predictions of epistasis. This is the most generalizable quantity to assess cross-modular interactions and is as mentioned in the introduction critical for constructing GP-maps. For this purpose, we considered high-throughput data on numerous mutants, with varying levels of detail regarding the mutant phenotypes, which we define as the information content. This information will determine the precision with which the mutant can be incorporated into the model.

Concretely, we restrict ourselves to epistasis between general mutants and Δ*bem1*, since we suspected that fitness differences in this ill background are exaggerated and hence more likely to have been picked up in literature. We used Bayesian analysis on the prevalence of epistasis signs to determine what degree of information on the general mutants add value to sign predictions. The general mutants were absorbed in the model in three different ways; either using the coarse information on the single deletion phenotype (deleterious or beneficial), or the mid-detail information on the single deletion phenotype (faster, slower, larger or smaller in G1), or the functional information (proteasomal, phospholipid or ribosomal). Within these three categories, there is a further subdivision into two sets, based on whether the model predicts positive or negative epistasis with Δ*bem1* (Fig. 5A).

**Fig 5.**
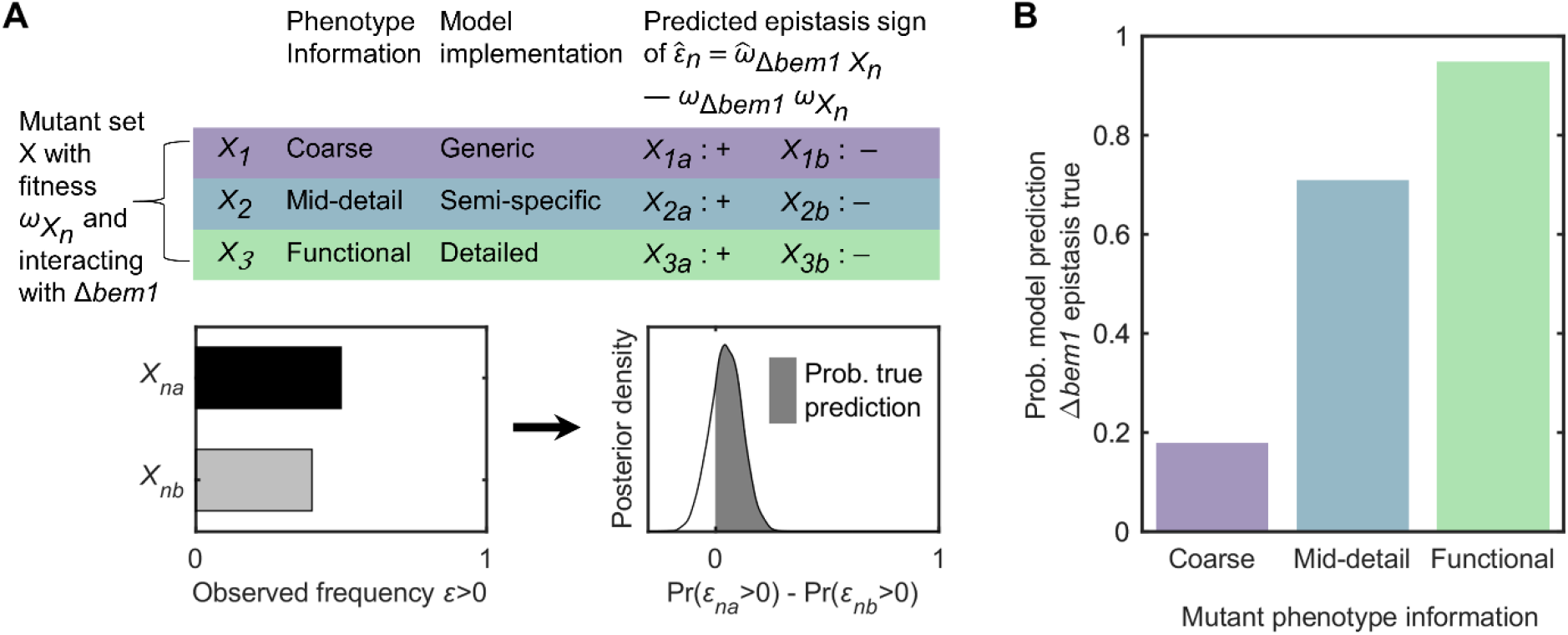
Adequate model predictions of epistasis rely on functional information concerning mutations. (A) Workflow for model prediction on epistasis *ε* of general mutants (fitness *ω*_*X*_) with Δ*bem1*. Mutants are divided into three categories and two subsets, depending on the specificity of the mutant phenotype and model implementation and the subsequent model prediction. For each category *n* and subset *a*/*b*, the beta posterior density of the observed positive epistasis fraction can be constructed (from a binomial likelihood an uniform prior). The probability of a true prediction is then defined as the area below the posterior density of the difference of sets *a* (prediction *ε*>0) and *b* (prediction *ε*<0). (B) Bars reflecting Bayes factors for the model hypothesis; the ratio between the odds that the model prediction is true and false.

Firstly, mutants of which the coarse information are incorporated through modifying the membrane area growth rates, concretely smaller and larger rates for deleterious and beneficial mutants respectively. As seen in Fig. 4, smaller rates reduce the deleterious effect of the Δ*bem1*, prompting the prediction that negative epistasis with Δ*bem1* is generally more prevalent for deleterious mutants than for beneficial mutants. The analysis shows no evidence that this statement is correct (only a 20% chance, Fig. 5B).

Analogously, integrating the mutants on mid-detail information implies changing *t*_*G1,min*_ (shorter when fast in G1, longer when slow) or *r*_*min*_ (lower when small in G1, higher when large). Mutants with shorter *t*_*G1,min*_ and lower *r*_*min*_ disproportionally benefit the Δ*bem1* which suffers most from Cdc42p dilution before the mesotype checkpoint. Therefore, the model prediction is that mutants fast or small in G1 have more negative epistasis with Δ*bem1* than mutants that are slow or large in G1. Still, the experimental evidence is not compelling (70% chance).

Finally, when incorporating the mutants using functional information, we lower *τ*_*h*_ (proteasomal), membrane growth rates (phospholipid) and mean burst size *p*_*b*_ (ribosomal). The former two, which mitigate the problematic lack of Cdc42p in the Δ*bem1* to some extent, should exhibit more negative epistasis than the latter one, which deteriorates the Δ*bem1* situation. There is strong positive evidence for this statement (using the rules-of-thumb on Bayesian odds ratios [44]), which is true with around 95% certainty. This displays the benefit of integrating mutants based on functional information.

## Discussion

We have constructed, verified, validated and applied a coarse-grained growth model encompassing the newly defined mesotype in order to describe phenotypes (subject to epistasis) from genotypes or predict these. When ample molecular information is present, as is the case for Bem1p and the GAPs, this strategy is quite successful to predict cell cycle times, given the largely good quantitative matches in Fig. 3A and C and qualitative match for the peculiar G1 time inversion for the Δ*bem1* compared to WT (Fig. 3B).

Additionally, the information content about the phenotypes, associated with mutated genes, required for predicting epistasis was assessed as it is a general hurdle for GP-map models. As the mutant is encapsulated in our model through more detailed phenotypes, the prediction quality increases accordingly. Typically, functional information is required to make meaningful epistasis sign predictions (Fig. 5), similar to the ontotype strategy [18]. This delimits the scope of this model.

Nevertheless, when only medium detail phenotypical information on the single deletion mutant (such as in the case with Nrp1p) is used, predictions can still be of decent quality (Fig. 3A). The efficacy of phenomenologically integrating Nrp1p into this model provided substance to the claim that this protein is mechanistically involved in shortening G1. Since obtaining near-complete information on the function of proteins is not within reach for most organisms, it is comforting that mildly positive results may be achieved with phenomenological information when building an otherwise biophysically justifiable bottom-up model.

Because the yeast polarity example shows the feasibility of our modelling strategy, we aim to provide a road-map to apply these to general genotype-phenotype maps (Fig. 6). The core functional component, in this case polarity, is modelled by justifiable coarse-graining, which results in the mesotype of the system. This mesotype in turn emerges from functional subunits [32], identifiable from the rigorous analysis of the underlying biophysics. Once multiple model systems (such as the PAR protein system in *Caenorhabditis elegans* [45]) have been described in this manner, it may be possible to construct a limited library of recurring subunits, making it easier to recognize these in other systems and construct the corresponding mesotype. In combination with a coarse-grained view of cell growth and noisy protein production, this completes the bottom-up (population) phenotype prediction process.

**Fig 6.**
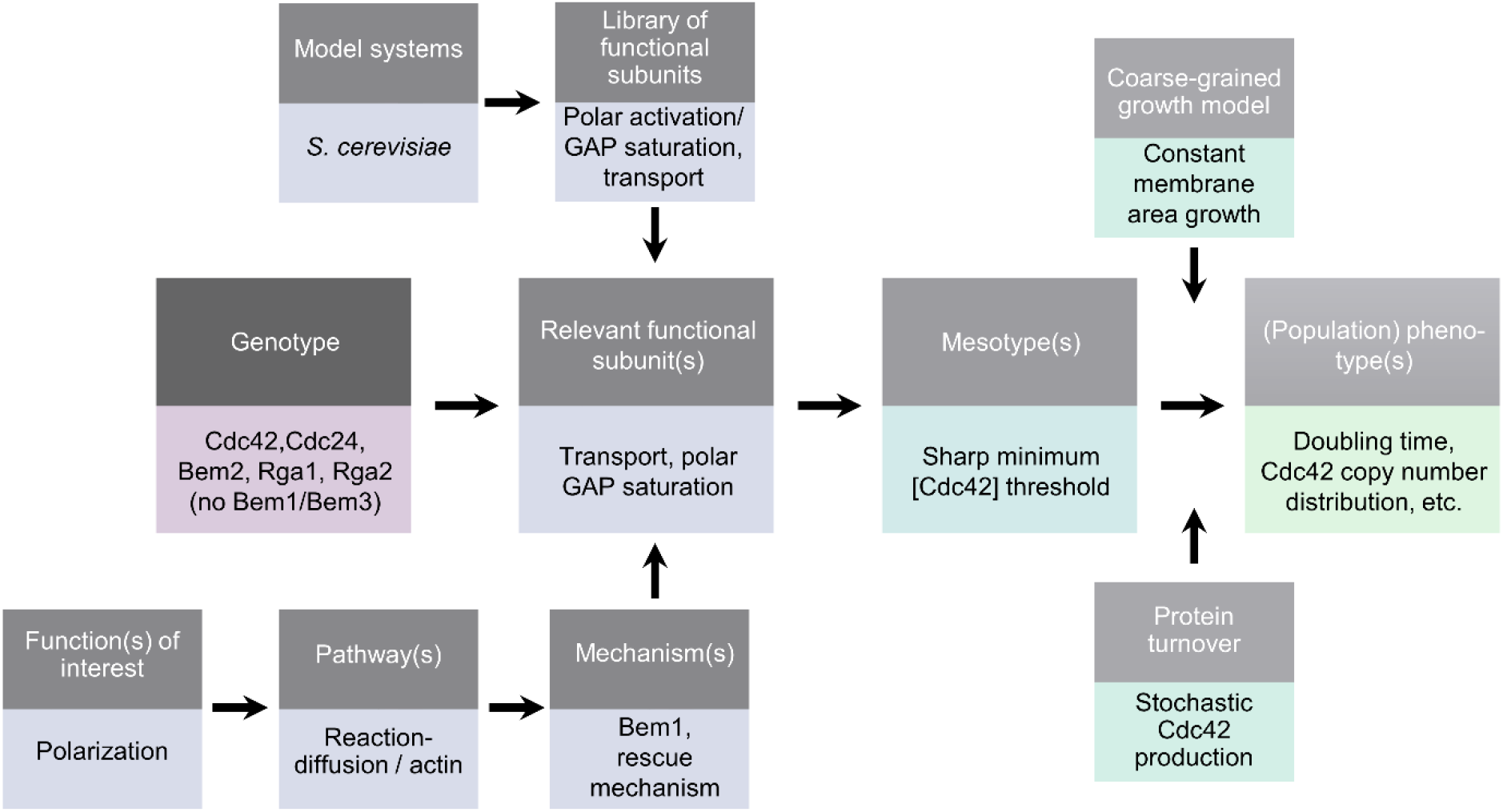
Proposed flow chart for phenotype predictions through intermediate levels. Bottom-up approach for phenotype prediction from genotype through mesotypes, which result from selecting the appropriate functional subunits. When more model systems are analyzed (e.g., polarization in *S. cerevisiae*, the min-system in *E. coli*, PAR-system in *C. elegans*), the toolbox from which to retrieve the relevant subunits expands. While it is possible to bypass the functional subunits and retrieve the mesotypes with rigorous numerical analysis of simulations of all protein components, the path displayed lends itself better to transfer knowledge of mesotypes to other systems. Bottom-half exemplifies the flow chart with the yeast polarity case.

Furthermore, the benefit of this approach is the tractable identification of evolutionary relevant quantities. For example, the GAP epistasis is accurately retrieved (Fig. 3A), and the prediction of the poor medium effect to reduce fitness differentials (Fig. 4) readily allows interpretation. The benefit of slower medium for the ill mutant Δ*bem1* fits the picture that haploinsufficiency in YPD is typically lifted in poorer medium [46], and opens up a distinct avenue for adaptation. Given that laboratory conditions are much more comfortable than the conditions under which historical evolution has taken and is taking place, the likelihood of fixation of a polarization network optimized on Bem1p or an rescue mechanism (as experimentally occurring in [26]) becomes much more similar than naively expected. Moreover, this insight is quantifiable, we show e.g., that merely slowing WT down by a factor of 2 reduces the relative fitness differential to 0.05. Given that *BEM1* has comparable characteristics to an essential gene, the evolvability of essential genes may be greater than anticipated.

## Materials and Methods

Model simulations were performed in MATLAB R2016a following a partial leap-like Gillespie algorithm [47] implementation (the G1 time until *r*<*r*_*min*_ and *t*_*1*_<*t*_*G1,min*_, *t*_*pol*_ and the time through S/G2/M are one leap each). The core function and example script to demonstrate the functionality are found in S1 Code and S2 Code respectively.

Model parameters are summarized in S2 Table. An initial population asynchronized across a bandwidth of 83 minutes (all cells with equal radii of 2.2 μm and without protein) is grown until a population size of >5 million, after which a subsample of 1000 cells is regrown to the same condition. Doubling times are the average of the last hundred moving window (size 201 min.) linear regressions on the log number of cells.

Model calibration was done by supplying expression burst parameters for Cdc42p inferred from flow cytometry. These were fine-tuned, together with area growth rates *C*_*1*_ and *C*_*2*_, to yield a mean protein copy number of around 8700 [41] at the optimal growth doubling time of 83 minutes (WT in YPD [26]). Fluorescence measurements of the required C*DC42pr-GFP-CDC42* strain and a non-fluorescent strain (from [48,49]) were performed using a BD FACScan flow cytometer. Cells were pregrown in YNB (Sigma) + CSM -Met (Formedium) + 2% dextrose (Sigma-Aldrich), diluted to an OD_600_ of 0.1 and measured after 15h.

Doubling times of [26] in Fig. 3A were fitted using the native *fminsearch* on a normalized score objective for varying [Cdc42]_min_ and manual inspection for setting *t*_*mut*_ (to 0.75) for the *nrp1* deletion. Interaction and phenotype data for Fig. 5 were obtained from BioGRID [50] and SGD [51] respectively (date of access March 2018).

## Acknowledgements

We thank Marit Smeets for flow cytometry measurements of both strains and Leila Iñigo de la Cruz for growth data on one of these. Additionally, we thank Fridtjof Brauns for careful reading of the manuscript and Sophie Tschirpke for help in optimizing figure formatting.

## Supporting Information

The flow cytometry data (see S1 Dataset) was acquired using FlowJo CE software with a BD FACScan and later analyzed with home-written code in MATLAB. A single gamma distribution was maximum likelihood fitted on the fluorescence intensity counts of strains (from [48,49]) with simply endogenous *CDC42* (‘background’) or *CDC42p-GFP-CDC42* at 2% dextrose (‘WT expression’). The WT expression distributions was analytically deconvolved for background counts using a gamma-sum approximation [52]. The average burst interval duration and average burst size result from these normalized distributions [39], a coarse doubling time estimate (200 min., processed as in [32]) for RWS1421 and the Cdc42p copy number estimate of 8700 from [41]. Calibration shows average burst sizes require a 20% reduction due to explicit inclusion of degradation in our model.

**S1 Fig.**
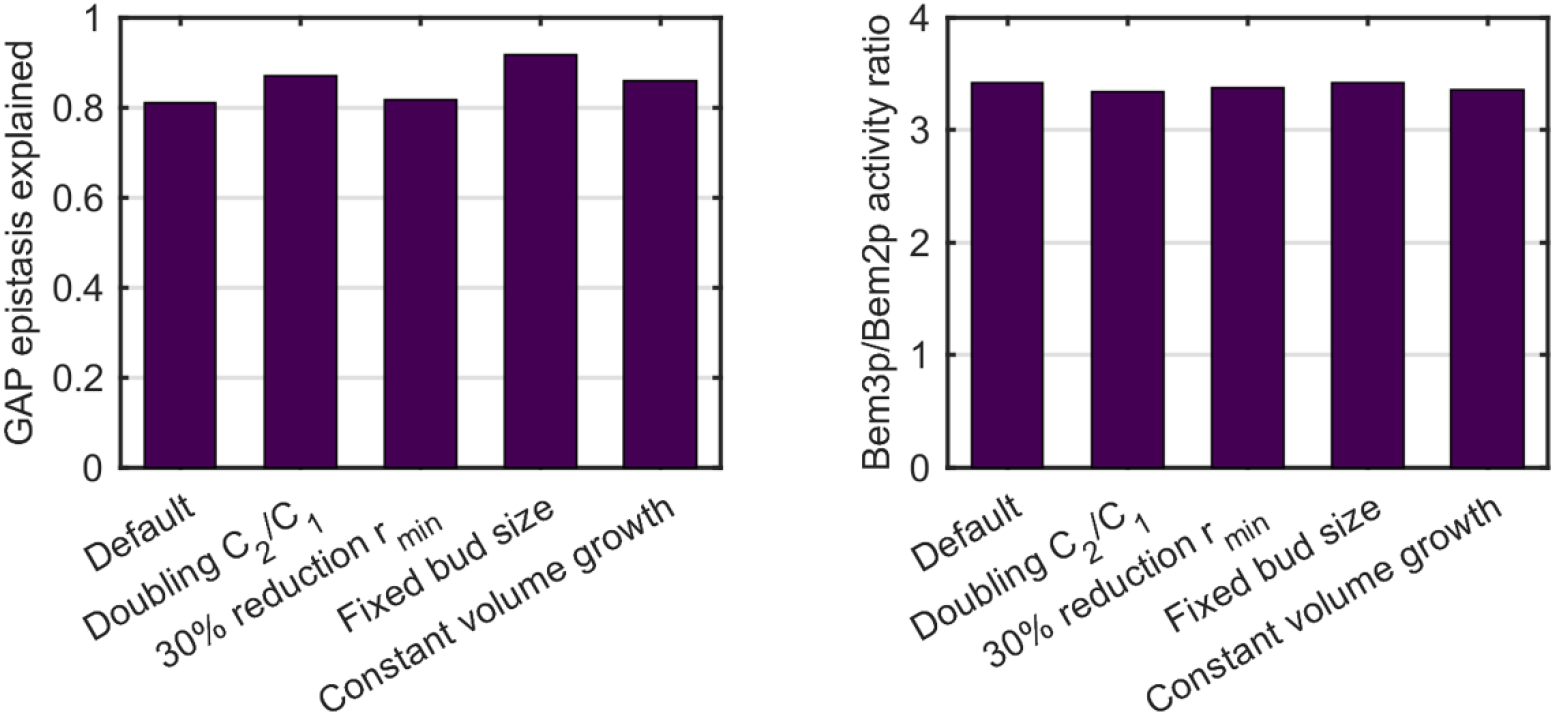
Negligible influence of model details on GAP epistasis. Calculated GAP activity given model fits and abundancies from [41] (left) and relative multiplicative epistasis (definition of [53]) for the GAPs in the Δ*bem1 NRP1* background for the growth model and four variations; doubling of membrane area growth rate *C*_*1*_, 30% reduction of *r*_*min*_, change of second checkpoint to a fixed bud size threshold of 1.8 μm, and constant cell volume instead of area expansion. WT membrane growth rates are recalibrated in each case to match 83 minutes.

**S2 Fig.**
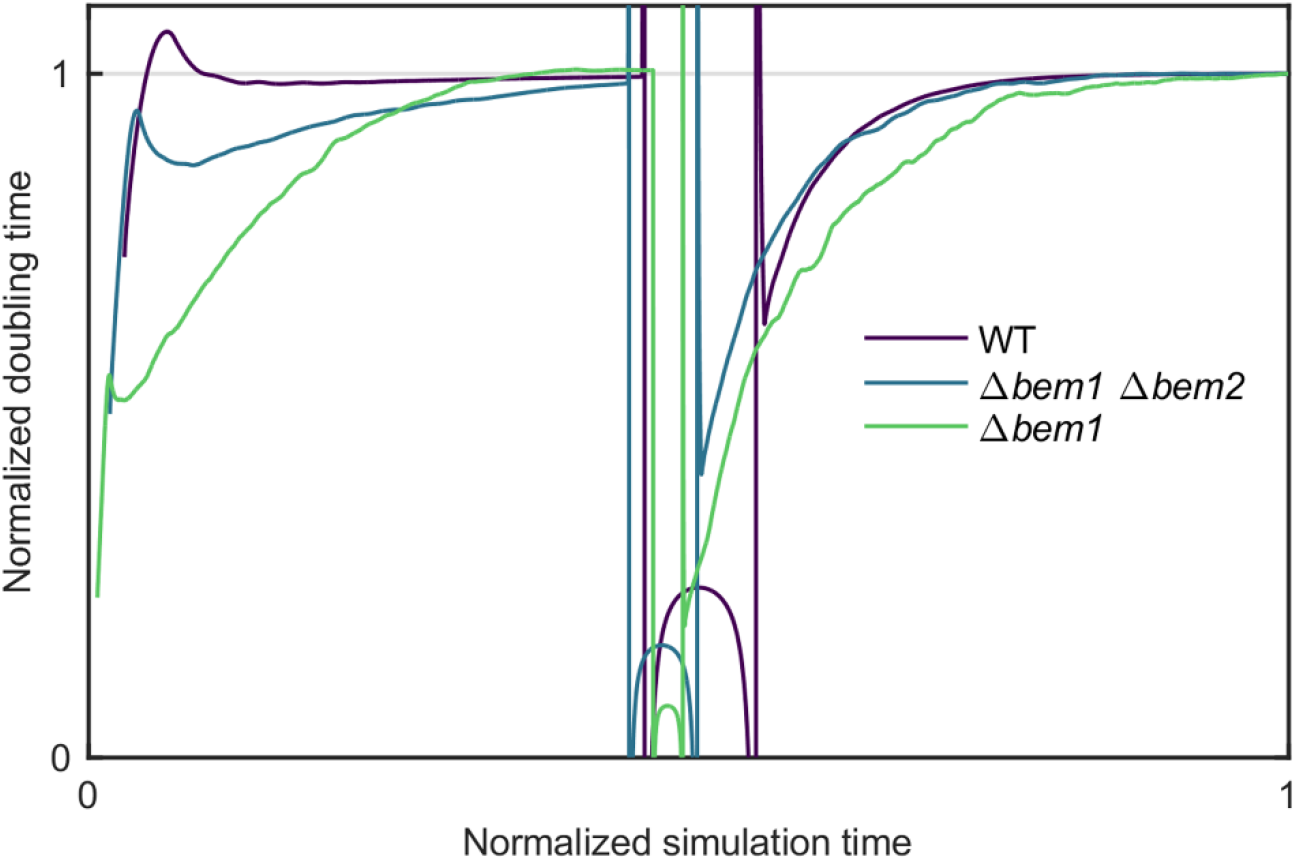
Convergence of simulated doubling times. Doubling times as function of simulation time for a fast (WT), medium (Δ*bem1* Δ*bem2*), and slow (Δ*bem1*) growing strain background. Doubling times and simulation times normalized to their respective final value. The dilution step midway temporarily causes unreliable estimates.

**S1 Table.**
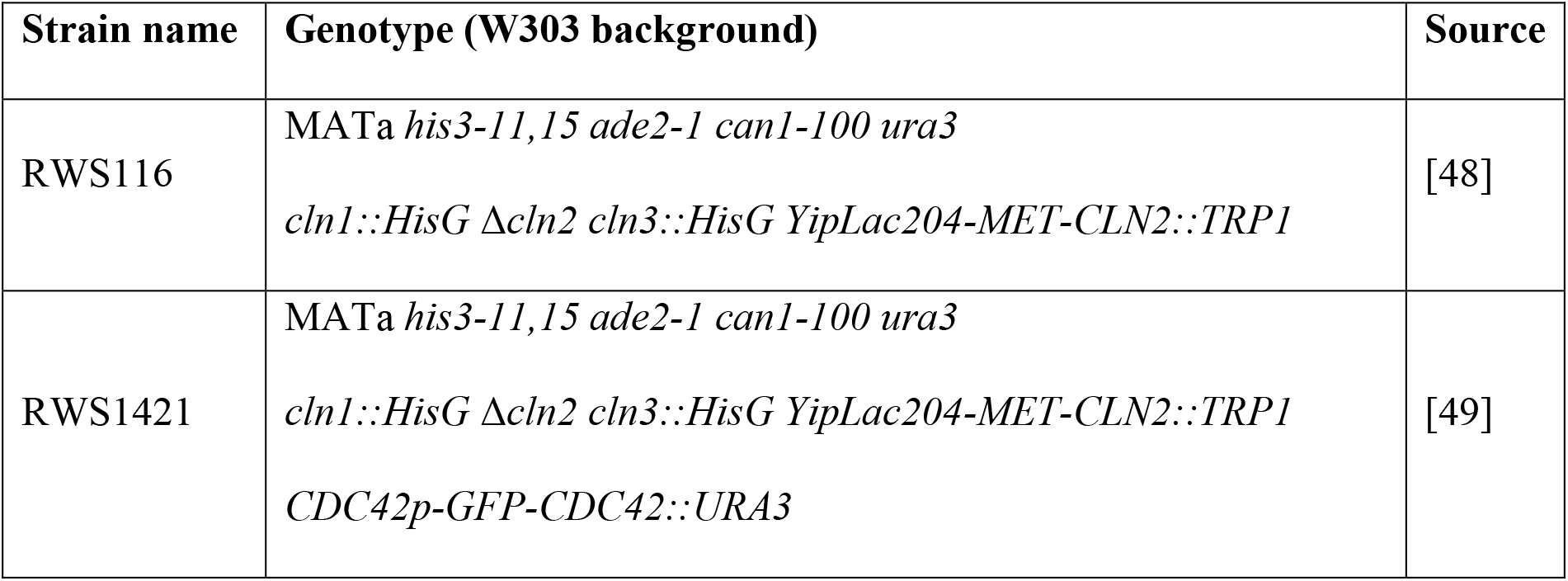
Strain list. Strains used in this study.

| Strain name | Genotype (W303 background) | Source |
| --- | --- | --- |
| RWS116 | MATa *his3-11,15 ade2-1 can1-100 ura3*<br>*cln1::HisG Δcln2 cln3::HisG YipLac204-MET-CLN2::TRP1* | [48] |
| RWS1421 | MATa *his3-11,15 ade2-1 can1-100 ura3*<br>*cln1::HisG Δcln2 cln3::HisG YipLac204-MET-CLN2::TRP1*<br>*CDC42p-GFP-CDC42::URA3* | [49] |

**S2 Table.** Growth model parameter list.

| Parameter | Symbol | Value<br>(background) | Source |
| --- | --- | --- | --- |
| Cdc42 concentration threshold | $[Cdc42]_{min}$ | 115 # proteins/$\mu m^3$<br>($\Delta bem1$)<br>4 # proteins /$\mu m^3$<br>($BEM1$)<br>-62% ($\Delta bem3$)<br>-34% ($\Delta bem2$) | This study, fitted |
| Minimum G1 time multiplier | $t_{mut}$ | 1 ($NRP1$)<br>0.75 ($\Delta nrp1$) | This study, fitted |
| Minimum polarization time | $t_{pol,min}$ | 5 min. | [54] |
| Maximum polarization time | $t_{pol,max}$ | 600 min. | To truncate computations for cells with extremely low GAP content |
| Polarization time scaling parameter | $\phi_{pol}$ | 25 ($\Delta bem1$)<br>500 ($BEM1$) | $\phi_{pol,BEM1} \gg 1$ and<br>$\phi_{pol,BEM1} / \phi_{pol,\Delta bem1} \cong$<br>$[Cdc42]_{min,\Delta bem1}$<br>/$[Cdc42]_{min,BEM1}$ for observed small excess Cdc42 across backgrounds (this study) |
| Minimum G1 time | $t_{G1,min}$ | 15.6 min. | [35] |
| Minimum radius to polarize | $r_{min}$ | 2 $\mu m$ | [35] |
| Maximum radius in G1 | $r_{max}$ | 6 $\mu m$ | To truncate computations for cells with very low Cdc42 content |
| Average Cdc42 expression burst interval time | $t_{b,WT}$ | 57 min. | This study, assuming theory from [39] |
| Average Cdc42 expression burst size | $p_{b,WT}$ | 4900 | This study, assuming theory from [39] and with calibration |
| Cdc42p half-life | $\tau_h$ | 474 min. | [38] |
| Bud/mother volume ratio checkpoint 2 | $c_r$ | 0.89 | Consistent with [35] |
| Ratio polarized/isotropic membrane area growth rates ($C_2/C_1$) | $c_p$ | 2.13 | Calibration |
| Isotropic membrane area growth rate | $C_1$ | $0.086 \mu\text{m}^2/\text{min.}$ | Analytical considerations for optimized WT (see S1 Text) |

**S1 Dataset. Flow cytometry and growth assay data.** Flow cytometry data (raw, processed and fitted) of strains used in this study, and OD_600_ measurements of one strain (with fits).

**S1 Code. Numerical implementation in MATLAB of the growth model in this study.**

**S2 Script. Test script in MATLAB calling the numerical model implementation of S1 Code.**

**S1 Text. Justification of membrane area growth rate value.** We assume an optimized WT such that at checkpoint 1, the minimum size requirement is typically met at the same time that the minimum G1 time requirement is met, and that subsequent polarization time is minimal. After G1, the (squared) mother radius is then (integrating the radius equation from 0 to *t*_*pol,min*_):

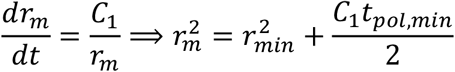

Thus, the bud radius is after next M-phase *r*_*b*_=0.89 *r*_*m*_. For self-consistentcy, this new cell must then expand to size *r*_*min*_ again at checkpoint 1, such that:

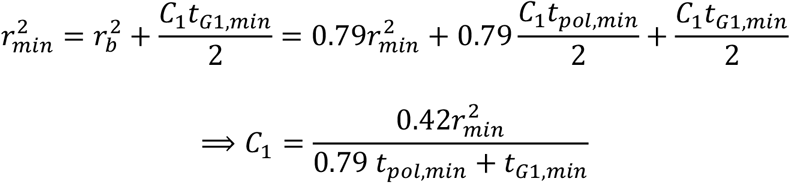

Using the values from S2 Table for *r*_*min*_, *t*_*pol,min*_, and *t*_*G1,min*_, this leads to *C*_*1*_=0.086 μm^2^/min.

